# A configural context signal simultaneously but separably drives positioning and orientation of hippocampal place fields

**DOI:** 10.1101/2023.05.28.542182

**Authors:** Han Yin Cheng, Dorothy W Overington, Kate J. Jeffery

## Abstract

Effective self-localization requires that the brain can resolve ambiguities in incoming sensory information arising from self-similarities (symmetries) in the environment structure. We investigated how place cells use environmental cues to resolve the ambiguity of a rotationally symmetric environment, by recording from hippocampal CA1 in rats exploring a ‘2-box’. This apparatus comprises two adjacent rectangular compartments, identical but with directionally opposed layouts (cue card at one end and central connecting doorway) and distinguished by their odor contexts (lemon vs. vanilla). Despite the structural and visual rotational symmetry of the boxes, no place cells rotated their place fields. The majority remapped their fields between boxes but some repeated them, maintaining a translational symmetry and thus adopting a relationship to the layout that was conditional on the odor. In general, the place field ensemble maintained a stable relationship to environment orientation as defined by the odors, but sometimes the whole ensemble rotated its firing *en bloc*, decoupling from the odor context cues. While the individual elements of these observations – odor remapping, place field repetition, ensemble rotation and decoupling from context – have been reported in isolation, the combination in the one experiment is incompletely explained within current models. We redress this by proposing a model in which odor cues enter into a three-way association with layout cues and head direction, creating a configural context signal that facilitates two separate processes: place field orientation, and place field positioning. This configuration can subsequently still function in the absence of one of its components, explaining the ensemble decoupling from odor. We speculate that these interactions occur in retrosplenial cortex, because it has previously been implicated in context processing, and all the relevant signals converge here.

## Introduction

Place cells fire in specific locations in an environment, forming the neural basis for the brain’s internal representation of space (‘cognitive map’; O’Keefe and Nadel, 1978). Place cells use environmental features to localize their firing (K.J. Jeffery, 2007), so how does the system distinguish environments that have repeated identical elements, and locate the animal within each one?

We have been exploring this issue using environments possessing an inherent symmetry, broken only by sensory cues of interest. Symmetry refers to a system’s property of invariance under a transformation. For a spatial environment, symmetries of relevance are translational, rotational and scale (Kate J Jeffery, 2023). If place cell firing patterns break these symmetries (respond differently in the environments), this indicates that these cues are used by the network, and suggests how: for orientation, positioning etc. The emerging picture so far is as follows. First, the cells distinguish identically shaped and oriented environments using context cues such as odor (Anderson & Jeffery, 2003): essentially breaking a translational symmetry, where odor is the dimension of translation. By contrast, they do not readily do this using path integration (movement-tracking; Spiers et al., 2015; Grieves et al., 2016). They do, however, use path integration to break rotational symmetry, enabling them to, for example, distinguish one end of a rectangular compartment from the other (K J Jeffery, Donnett, Burgess, & O’Keefe, 1997; Taube & Burton, 1995). They also readily use local (mainly visual) cues such as landmarks, to do this (Muller & Kubie, 1987). This rotational symmetry-breaking is thought to be mediated primarily by head direction (HD) cells (Harland et al., 2017; Taube, Muller, & JB, 1990).

HD cells are central to the organization of these firing patterns, so we recently asked the question of how the HD system itself solves this same problem of resolving environmental symmetries. We found to our surprise that a subset of cells in rat dysgranular retrosplenial cortex (dRSC), which outwardly resemble classic HD cells, are responsive to local but *not* global rotational symmetry-breaking cues (Jacob et al., 2017; Zhang, Grieves, & Jeffery, 2022). This was found by recording in the ‘2-box’, which comprises two connected rectangular compartments in which the visual layouts are 180 degrees reversed from each other. This arrangement creates a twofold global rotational symmetry for the whole apparatus. Additionally, if the door and cue cards are ignored there is also a translational symmetry and a local onefold rotational symmetry, arising from the rectangular geometry (Figure 1). We found that as the animal crossed from one box to another through the doorway in the central dividing wall, some cells ignored the global symmetry-breaking cues and rotated their directional firing to remain aligned with the local visual layouts. We proposed that these layout-following ‘multidirectional’ cells represent a link in the chain between local visual scene-processing and global head-direction orientation (Page & Jeffery, 2018). These experiments also revealed that ordinary HD cells can break the symmetry of the two-compartment layout by using the odors in the two compartments, even though odor is not in and of itself a precise directional cue.

**Figure 1.**
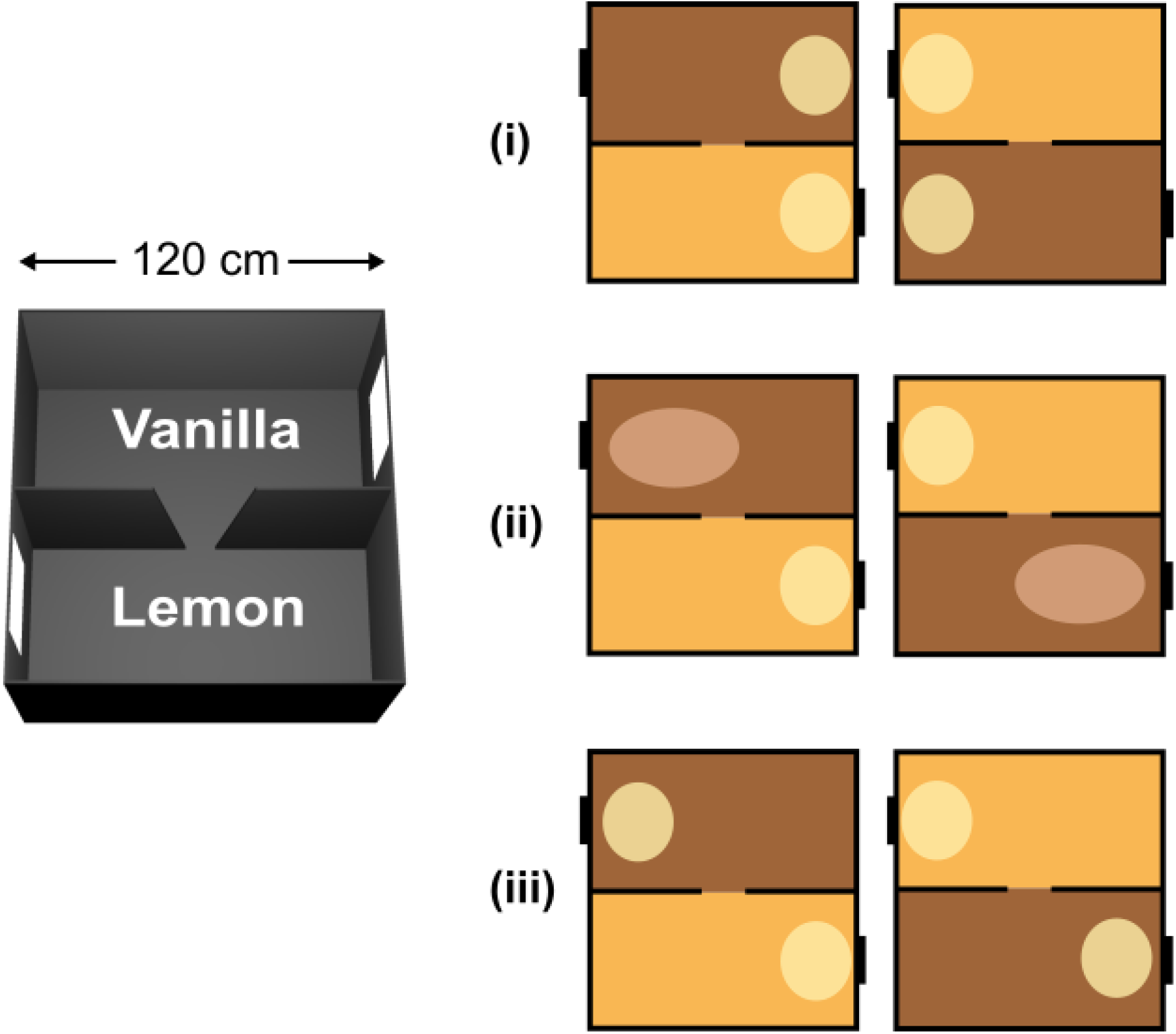
Multiple symmetries in the two-compartment context box. Left: The two-compartment context box comprises two visually identical compartments, painted grey and scented with either lemon or vanilla. The local symmetry of each sub-compartment is broken by the asymmetrically placed visual cues (cue card, plus the doorway). The global symmetry is broken by the relative positions of the two odors. Right: Three types of symmetry-breaking that could be detected by place cells (see text). (i) The place fields (ovals) could be identical in both compartments but both located at the same end, which is a translational symmetry. This would mean they do not use local odor cues nor local visual asymmetry, but are oriented by the global directional cues. (ii) The cells might remap between compartments. Here, symmetry could be locally broken by the odors (inducing contextual remapping) or by the directional cues, in which the system detects that the cue cards have different relationships to the global directional orientation. (iii) The cells might show a rotationally symmetric pattern, with fields in the same local location in the two sub-compartments, ignoring both local odor context and the global odor-referenced directional cues.

The question arises as to whether all this information is passed to place cells. Thus, the present study used this same 2-box environment with its multiple symmetries to investigate the spatial organization of place fields. Given our finding that the retrosplenial HD signal is not unitary, but rather has cells with differential sensitivity to the global vs. local directional cues, we wondered what place cells would do. Would they (see Figure 1):

i. Repeat their fields between the two compartments, showing translational invariance, and sensitivity to global but not local rotational symmetry-breaking
ii. Change their firing (remap), showing sensitivity to either translational or rotational symmetry-breaking cues (or both), or
iii. Rotate, showing sensitivity to local rotational symmetry-breaking cues (the visual layout) but insensitivity to both the translational *and* the global rotational symmetry-breaking cues.

Furthermore, would their responses be global (across the whole population) or partial (affecting some cells only)?

We found here that only (i) and (ii) held true; place cells repeated or remapped their fields but never rotated them. In terms of complete vs. partial responding, place cells showed complete (ensemble) responses to global rotational symmetry-breaking cues, but partial responses to the local cues. Furthermore, sometimes they collectively and spontaneously broke the global symmetry independently of the odor cues, indicating more than one way to signal global orientation.

This behavior is not readily explained by current models of place field orientation and positioning. We therefore propose a new model, adapted from an existing ‘contextual gating’ model, that could explain these findings and map them to known circuit properties.

## Materials and methods

### Subjects

Five Lister Hooded rats were used for place cell recordings. All animals except one were naïve to the experimental apparatus prior to any recordings. The remaining animal had been exposed to the experimental apparatus as part of a behavioral experiment before entering the current experiment.

The animals were housed individually and maintained on a 12hr/12hr light/dark cycle. A week after surgery and throughout the whole experimental period, all animals were food restricted and their body weight maintained at 90% of their free-feeding body weight. All procedures were done in accordance with the restrictions and provisions listed in the Animals (Scientific Procedures) Act (United Kingdom, 1986) and European Communities Council Directive of November 24,1986 (86/609/EEC)).

### Microdrive implantation

Axona microdrives (Axona Ltd, UK) containing four drivable bundles of tetrodes were prepared for implantation in rats. Each tetrode was made from four 25µm polyimide coated 90% platinum-10% iridium wires (California Fine Wire, Grover Beach, CA) twisted together and threaded through a 21G stainless steel cannula (Cooper’s Needle Works Ltd, UK). Gold plating was used to reduce impedance to within the range of 180-300 kΩ (with a target impedance of 200 kΩ).

Standard procedures were used for stereotaxic implantation of microdrives into the cortex overlying the hippocampus (AP: −4.0, ML: ± 2.5). Carpofen (0.1ml of 10% v/v per 100g animal weight; Pfizer Ltd, UK) was given pre-surgically while Metacam was given post-surgically to reduce pain and inflammation.

### Experimental apparatus

The animals were screened in a separate room distinct from the recording room. The screening room contained a square open box environment (120cm x 120cm) lifted off the floors by four stools, one in each corner. Strips of polypropylene sheeting were mounted onto the box wall to discourage the animals from climbing on or over the edge. Distal cues were available in the screening room.

The recording room contained the recording environment (2-box) surrounded by black floor-ceiling curtains, thus depriving animals of distal visual cues. The 2-box was constructed from wood, painted black and lined with polypropylene sheets for the walls, and vinyl sheet for the floor. A white cue card (A4-sized; 21cm x 29.7cm) was mounted on one short end of each compartment, ‘North’ for one and ‘South’ for the other. The floors were scented at the start of every trial by wiping the vinyl flooring with a diluted (∼50%) solution of either vanilla or lemon odors (Dr Oetker, Germany). Four halogen lamps were mounted onto the ceiling near each corner of the environment to provide symmetrical lighting, and white noise was played on a portable speaker mounted on the ceiling to mask extra-maze directional auditory cues. During recordings, the room lights were switched off and the halogen lights switched on.

### Recordings and experimental protocol

A week following surgery, the animals began screening for place cells in the screening room. For screening and recording, the connector on the microdrive was connected to a headstage that in turn was connected via a thin flexible cable to the dacqUSB pre-amplifier and recording system (Axona Ltd, UK). The signals acquired from the tetrodes were digitized (48 kHz), amplified 10000-20000 times and high-pass filtered to detect neuronal spikes. Meanwhile, local field potential signals were amplified 2000-3000 times and low-pass filtered. The positions of the animals were tracked via a single LED on the headstage via a ceiling-mounted camera. If place cells were observed, the animals were transferred to the recording room in a black transport box after which the recording protocol commenced.

Rats were always started in the same compartment, in order to generate an initial perception that the combination of box scent and box location is stable: an effect of entry location on directional initialization having been established in an early experiment by Sharp et al. (1990). Rats were not actively disoriented before the first trial, but were mildly disoriented in a gently rotated dark box between trials. Rice pops (Waitrose, UK) were thrown into the box to encourage the animal to explore while neural recordings were conducted. In two animals (R590 and R609), vanilla flavored rice was thrown into the vanilla compartment and non-flavored rice was used in the lemon compartment, to maximize the possibility for context discrimination. A single recording session consisted of five trials in sequence (Figure 2A): (i) baseline trial, (ii) rotation trial, (iii)-(iv) two single compartment trials, and (v) a second baseline trial. The rotation trial (90° or 180°) was to ensure that the cells were using the box cues and not uncontrolled distal cues, and the second baseline trial was to check the stability of the place cells. All two-compartment trials ranged from 15 to 25 min while the single compartment trials lasted for 10 min. In between trials, the animals were removed from the environment, placed back into black transport box and disoriented. The environment was cleaned of any urine or feces, the odors reapplied, and the box rotated to the appropriate configuration. At the end of a recording session, the screw on the microdrive was turned one-eighth of a turn (∼25µm) to find new cells for the next recording session.

**Figure 2.**
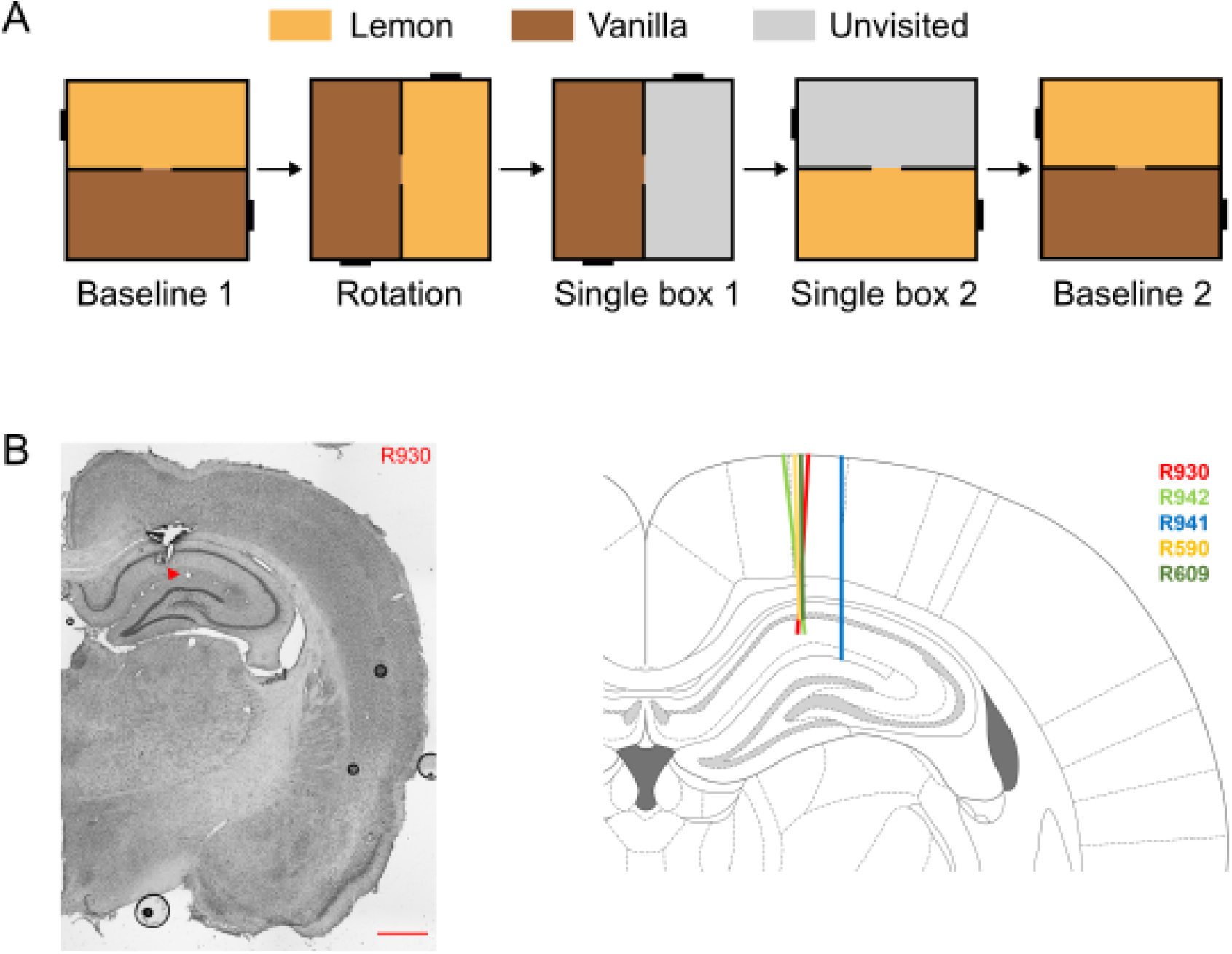
recording protocol in the two-compartment context box. (A) The experimental protocol showing the 5-trial sequence (door was closed for trials 3 and 4). (B) Recording locations, showing an example photomicrograph, and a schematic of the sites for the 5 animals.

Data were analyzed as follows.

### Behavior analysis

To examine for any bias in the animals’ preference for either compartment, a preference index was calculated as follows:

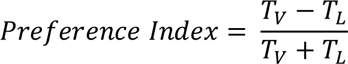

where T_V_ and T_L_ are time the animals spent in the vanilla and lemon compartments, respectively. PI values > 0 indicate a preference for the vanilla compartment while a PI < 0 indicate a preference for the lemon compartment. A value of 0 will indicate no preference for either compartment. The preferences for odors were only examined for two-compartment trials.

### Spike sorting

After recording, neuronal spikes were sorted using graphical cluster-cutting software Tint (Axona, UK) with the help of clustering algorithm KlustaKwik 3.0 (Kadir, Goodman, & Harris, 2014). Recording data obtained across trials in the same session were loaded into Tint together for clustering. The resulting clusters were manually refined by examining their waveforms and cross-correlograms, and merging them if they occupied similar spaces in the cluster space and did not spike within the 2 ms refractory period. The sorted spikes were further analyzed offline in Matlab (MathWorks).

### Place cell identification

Data from potential single units were preprocessed and analyzed in Matlab 2019a/b. First, positional data and spikes were speed-filtered. Spikes from epochs with an instantaneous speed < 5 cm/s were removed to exclude non-local spiking (replay events). Positional data and spikes with instantaneous speed > 100cm/s were considered to be tracking errors and removed.

To ensure good unit isolation, any units with more than 0.2% of spikes falling within an inter-spikes interval < 2ms and mean firing rate < 0.1Hz in both baseline trials were excluded. Units were considered to be putative pyramidal cells if the waveform peak to trough time > 250 µs and their mean firing rate was between 0.1 and 5 Hz in both baseline trials.

After identification of putative pyramidal cells, rate maps were constructed to examine for spatial firing. Rate maps were generated in the same manner as described in Leutgeb et al., (2005), but with slightly different parameters. Positional data were divided into 3×3 cm bins, and spikes falling within each bin were accumulated accordingly into spike maps, and position points similarly as dwell maps. Positions and spikes were smoothed separately with a Gaussian kernel (smoothing factor = 5 cm) centered on the bin. Any bins > 9 cm distant from the current bin were given a weight of 0. Bins in which the animals spent less than 0.1s were designated as unvisited. The smoothed rate maps were then generated by dividing the smoothed spike maps by the smoothed dwell maps.

Spatial firing was then examined using the smoothed rate maps. A place field was defined as region of at least 16 contiguous pixels with firing rates > 40% of the peak firing rate and peak firing of > 1Hz. Putative pyramidal cells with fields that had summed pixels > 50% of the dwell map pixels were considered ‘messy’ and excluded from further analysis.

To further remove cells with non-spatial firing, we calculated the spatial information content of each cell (Skaggs, McNaughton, Gothard, & Markus, 1992) and compared that to a shuffled null distribution. The shuffled null distribution is generated by circularly time shifting the spikes by a random time interval between 20s and the duration of the session minus 20s, for each round of the shuffle. This maintains the temporal relationship of the spikes but dissociates the spikes from the animal’s position. For each round of the shuffle, we generated a rate map based on the new spike positions and calculated the spatial information content. This process was repeated 100x for each cell, and the cell was considered to be spatially modulated (i.e. having a place field) if the cell’s spatial information content > 95^th^ percentile of the shuffled null distribution.

For visualization purposes, spikes were plotted on positional data for each trial to create a spike plot. Paths of the animals in the lemon compartment were plotted orange, and in the vanilla compartment in brown. Spikes of neurons from the corresponding time point were overlaid as red dots.

### Null distributions

To determine the chance level for which a place field is in a specific location between two correlation comparisons, null distributions were generated. To do this, randomly selected pairs of rate maps from the entire pool were correlated, and this procedure repeated 10,000 times. The 95^th^ percentile of the distribution of the resulting values was used as a threshold for the probability a place cell could have a place field in the exact same location by chance. Correlation values above threshold indicate similar rate maps while correlation values at or below threshold suggest dissimilar rate maps and/or global remapping.

### Rate maps correlation

To examine how similar or different spatial firing was across trials or compartments, Pearson’s correlations on the smoothed rate maps were carried out between two trials or two compartments respectively. Before correlation, all rate maps were rotated to the appropriate orientation (either same orientation or 180° rotation depending on the question to be answered). If the rate maps were found to be of different sizes (due to unvisited regions), the smaller rate map was padded to the size of the larger rate map with NaN in all dimensions. The equivalent spatial bins between the two rate maps were then correlated on a bin-by-bin basis. The threshold correlation value for what constituted a similar spatial map depends on the null distributions generated for each type of comparison. For the entire 2-box taken as a whole, the null distribution threshold was 0.37, while for each sub-compartment the threshold was 0.45.

### Duplicate cell detection/removal

To avoid over- or under-estimating the proportion of place cells that could display local or global encoding, place cells that were recorded over days were identified systematically using information about the waveform and the place fields. This method assumed that place cells in the two-compartment context box are stable and do not remap over days.

Place cells that displayed similar spatial firing were first identified by correlating the rate maps of all cells recorded from the same tetrode between consecutive recording sessions. Cell pairs with spatial correlation > 0.375 were then examined for similarities in waveforms using Tolias distances as a measure for waveform similarities (Duvelle et al., 2019; Tolias et al., 2007). Specifically, Tolias distances can be described using two separate metrics – Tolias distance 1 and Tolias distance 2 respectively. Tolias distance 1 captures the differences in waveform shapes and can be calculated by scaling one of the two waveforms to be compared by a factor α. After finding the best α fit for each channel, Tolias distance 1 can then be calculated by looking at the normalized Euclidean distance between the two waveforms. Tolias distance 2 examines the differences in amplitudes across the four channels. If the same cell was recorded over multiple days with minimal changes in waveforms, both Tolias distances will return small values.

We next sought to determine an objective threshold that would indicate if the two waveforms are similar or different. To do so, we fitted a linear discriminant analysis classifier using the Tolias distances calculated from experimentally defined *same cells* or *different cells*. *Same cells* were defined as the Tolias distances obtained by comparing the waveforms of cells determined to be the same (via spike sorting) across the two different two-compartment trials (baseline 1, rotation and baseline 2) within a recording session. Meanwhile, *different cells* were defined as the Tolias distances obtained by comparing the waveforms of cells recorded in the very first and the very last session of an animals. Separate linear discriminant analysis classifiers were generated for each animal under the assumption that each animal (and each microdrive) has its own set of electrical noise during recording.

Tolias distances were then calculated for cells recorded on the same tetrode on consecutive sessions, and the posterior probability of cell pairs having similar waveform (‘*same cell*’) predicted from the trained classifiers. Spatially similar cell pairs were considered to be the same if the trained classifier predicted a posterior probability > 0.5. Occasionally, two cells recorded in the first session mapped to a single cell in the next session (‘convergence’), or alternatively a single cell in the first session mapped to two different cells in the subsequent session (‘divergence’). In these cases, If the difference in the posterior probability between the two cell pairs exceeded 0.1, the cell pair with the higher posterior probability was considered to be the same cell. Alternatively, if the difference in the posterior probability was less than 0.1, the cell pair with the higher spatial correlation was considered to be the same cell.

Using this method, 104 cells were found to have been recorded in multiple sessions (2 sessions n = 70; 3-8 sessions n = 34). We removed these cells from every session except the first. In addition, we removed place cells that did not exhibit cue control (R < 0.37 between rotation trials) or were unstable (R < 0.37 across two baseline trials).

### Histology

At the end of the experiments, the animals were deeply anaesthetized with isoflurane (3% isoflurane with an O_2_ flowrate of 3L/min) and terminated via overdosing with sodium pentobarbital. The animals were then perfused transcardially with saline (0.9% sodium chloride solution) and 10% formalin. The tetrodes of the microdrives were then raised to the highest possible level so that aberrant tracks would not be made when the brains were extracted. The brains of the animals were then extracted and post-fixed in 4% paraformaldehyde for one or two days. They were then cryo-protected in 30% sucrose solution for approximately two days, before being sliced coronally into 40μm slices using a cryostat (Leica CM1850 UV). These slices were then mounted onto glass slides and Nissl-stained by immersing the slides in cresyl violet solution, washed in distilled water, dehydrated in increasing concentration of ethanol before being cleared in Histo-Clear (National Diagnostics, Scientific Laboratory Supplies). The slides were then cover-slipped using DPX mounting media (Sigma Aldrich).

All images were captured using a Leica DMi8 microscope under a bright-field setting. The images were acquired with a gain of 2.0 and an exposure time of 45ms. Whole-slice images were obtained by generating tile-scanned images which were stitched together with smoothed edges at 25x magnification.

## Results

The protocol for the experiment is shown in Figure 2A. All tetrodes were situated in the CA1 area of the hippocampus (Figure 2B). Five rats were recorded for 37 sessions of five trials yielding 437 unique place cells in the final analysis.

### Comparison of behavioral dwell time and place cell activity between compartments

First, in order to rule out potential effects of the odors, which may affect how place cells represent the environment (Mamad et al., 2017), we examined the preference of the rats for spending time in one compartment vs. the other and expressed it as a preference index (PI; see materials and methods, behavior analysis). We found a slight preference across all animals for the vanilla compartment (Figure 3A; one sample t-test t(110) = 3.59, p < 0.001). This declined over trials: a repeated-measures ANOVA found a main effect of trials [F(2,34) = 6.55, p = 0.002] with a significantly greater preference on the first trial than subsequently (baseline 1 vs rotation trials: t(36) = 3.08, p = 0.004; baseline 1 vs baseline 2 trials: t(36) = 3.33, p = 0.002) while the latter two trials did not differ significantly (t(36) = 0.50, p = 0.62). Thus, the vanilla preference slowly diminished over the trials such that by the rotation trial, the animals displayed no preference and explored both compartments equally.

**Figure 3.**
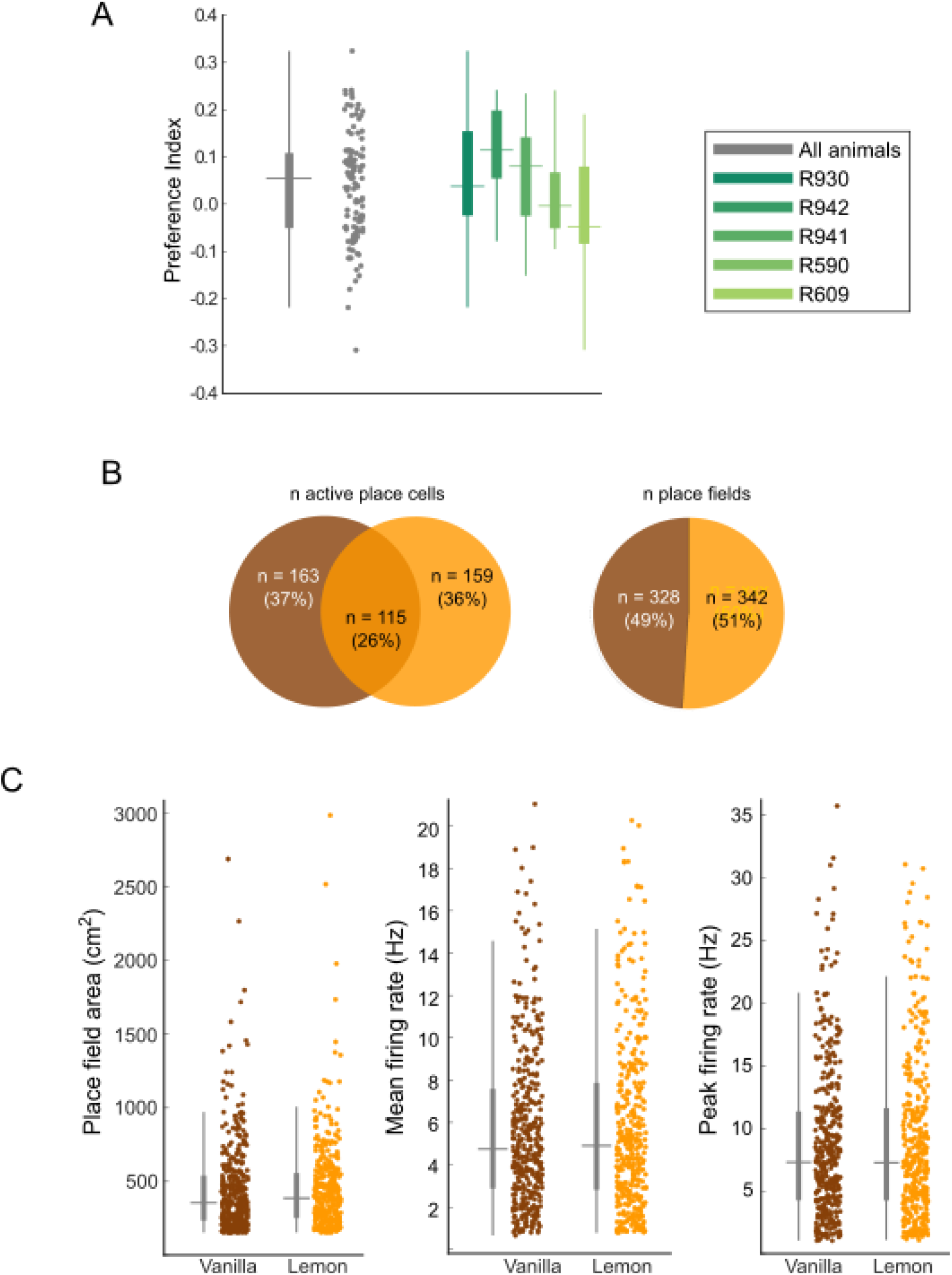
Comparison of basic firing properties in the two contexts. (A) Preference index of all trials (grey) and also averaged across trials for individual animals (colored bars showing mean, s.e. and inter-quartile range) showing mild preference for the vanilla compartment. (B) Comparison of the number of active cells (left) and place fields (right) in the two boxes showing no preference of the cells for encoding one vs. the other box. (C) Place field firing statistics did not differ between the two compartments.

We next compared the properties of CA1 place cells between the two boxes in Baseline trial 1, when this preference was the most apparent. Place cell activity did not differ between the boxes (Figure 3B) either in the number of active cells (*X^2^*(1, N = 322) = 0.075, *p* = 0.78) or the number of place fields (*X^2^*(1, N = 670) = 0.59, *p* = 0.44). There were also no differences in firing statistics (Figure 3C; Area: *t* test *t*(668) = −0.22, *p* = 0.83; Peak firing rate: *t*(668) = 0.14, *p* = 0.89; Mean firing rate: *t*(668) = 0.037, *p* = 0.97). Thus, overall there was little to distinguish the representations of the two boxes other than firing field locations, as analyzed below.

### Some place cells repeated their fields, most remapped and none rotated independently

We then looked at the spatial pattern of place cell firing in the two boxes, correlating the rate maps on a pixel-by-pixel basis under either translational or rotational transformation, or between the same box in different trials. The remapping correlation threshold was set to 0.45 based on the 95^th^ percentile of a shuffle (see Methods), and we evaluated the firing patterns based on the three predictions outlined in Figure 1. Note that sometimes the whole firing pattern rotated 180 degrees independently of the box (examined in more detail below) and so we corrected for this before comparing the firing patterns, as described in the Methods.

A small but substantial proportion of place cells (∼13.3%) repeated place fields between the boxes (see Figure 4A for examples), with average translational correlations of 0.66 ± 0.02 and rotational correlations of −0.12 ± 0.02. This proportion was significantly greater than chance (*X^2^*(1, N = 437) = 62.03, *p* < 0.001), and suggests spatial encoding independent of the local visual environment. The number of place cells with repeating place fields did not appear to change as a function of experience, and they could be recorded throughout the recording sessions.

**Figure 4.**
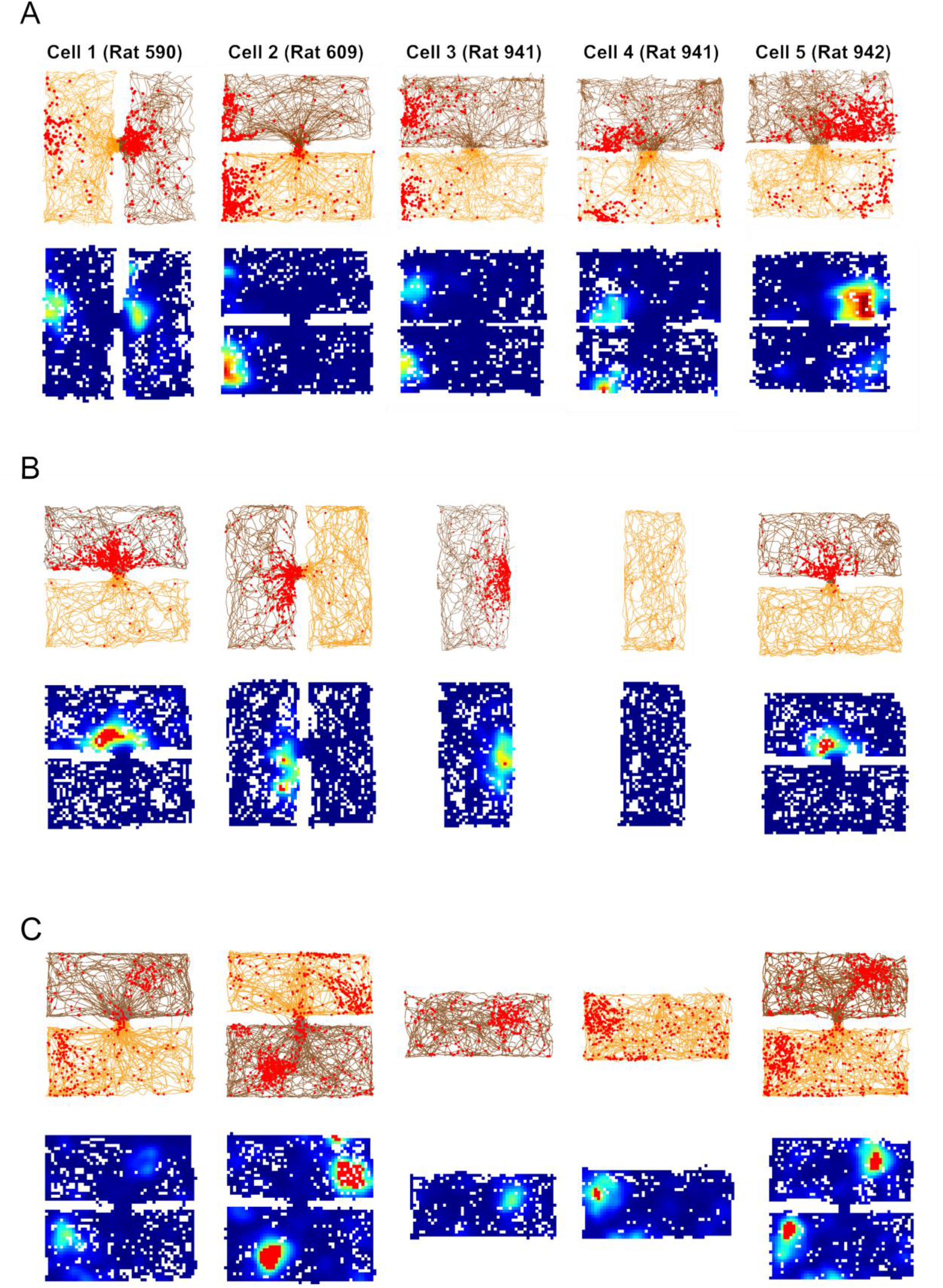
Examples of different types of compartment encoding. (A) Plots of five cells with repeating fields. Note that the firing fields are located at the same end of the box in a global reference frame even though both the odors and the local visual layout are different. (B) A cell that distinguished the compartments (remapped) by firing in only one of them, maintaining a constant within-compartment position. (C) A cell that remapped by shifting its firing location depending on the box identity. This cells thus receives two sets of positional inputs, which are selected (gated) by the odor context.

The majority of place cells (83.5%) had fields that remapped between the boxes (Figure 4B and C), with a low correlation for both translational and rotational comparisons (translational correlation of 0.041 ± 0.009 and a rotational correlation of −0.004 ± 0.009), but a high correlation for repeated trials in the same box (0.68 ± 0.009), which is greater than the 2-box remapping threshold of 0.37, indicating stable recording and stable encoding.

A small population of cells (∼3%) had rotated place fields, but visual inspection showed these to be spurious correlations from cells having very few spikes in one of the boxes. One cell fired exactly in the doorway and so had high correlations in both conditions: this cell was removed from the analysis. Figure 5 A and B summarize the proportion for each type of encoding.

**Figure 5.**
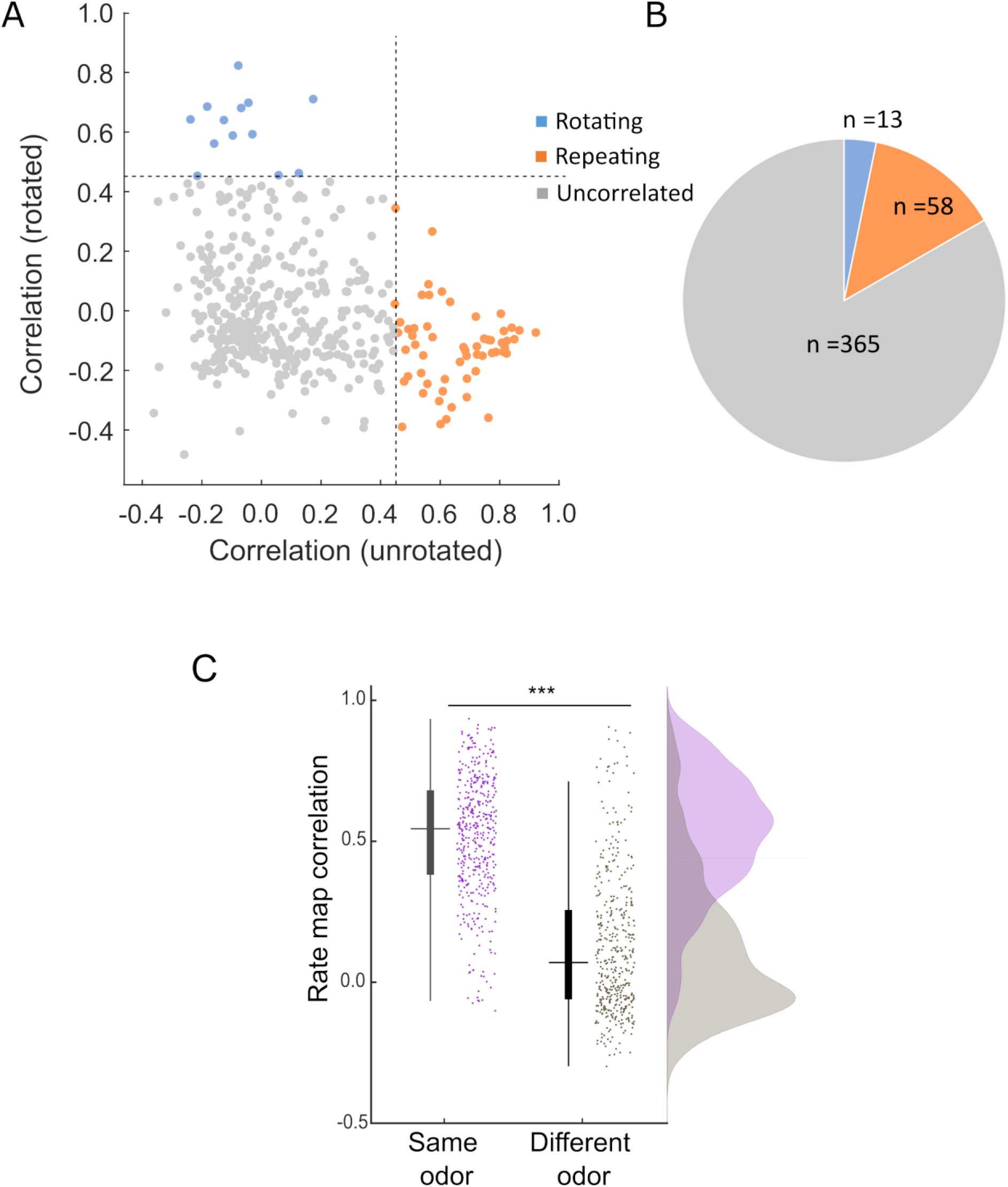
Quantification of remapping. (A) Scatterplot of correlations for all the place cells between the two boxes, either plotted in their original orientations, or with one of the boxes rotated. If fields repeat then correlations with the boxes in their original orientations are high, and in the rotated orientations low (orange data points). If fields rotate 180 degrees between the boxes then the reverse is true (blue data points). Cells with low correlations in either orientation had remapped (grey data points). (B) Pie chart of the data in (B) to show the predominance of remapping, and very small incidence of rotation. (C) Quantification of remapping across the odors vs. repeated trials in the same odor (each box in the rotation trial compared with the first single-compartment trial). All place cells were included in the analysis. Bars and lines show mean, s.e. and interquartile range; dots show individual cells. The low mean correlation for the different-odor comparison shows significant discrimination of the compartments based on their odor and/or relative location.

We examined whether place cells generated their box-specific firing patterns even in single boxes when the central door was closed. To do so, we aligned the boxes according to the visual scene and then correlated the immediately preceding rotation 2-box trial maps with those from the single-box closed-door trials. If place cells can distinguish the compartments during the single-compartment trials based purely on odors (i.e. not dependent on self-motion cues from moving across the compartment), same-odor maps should have significantly higher correlations than different-odor maps. This was indeed the case (Figure 5C; Two-tailed Wilcoxon signed rank test, Z = 17.46, *p* < 0.001). This indicates that place cells can use odor to recognize a single compartment, and self-motion cues across the two compartment are not essential for retrieving the correct maps.

### Place cells sometimes rotated their fields en bloc independently of odor

As mentioned earlier, when we examined the place cell maps within a recording session, we saw that sometimes place cells rotated their entire 2-box firing patterns 180° relative to the odors from one trial to the next (Figure 6A). We call this phenomenon ‘odor-switching’.

**Figure 6.**
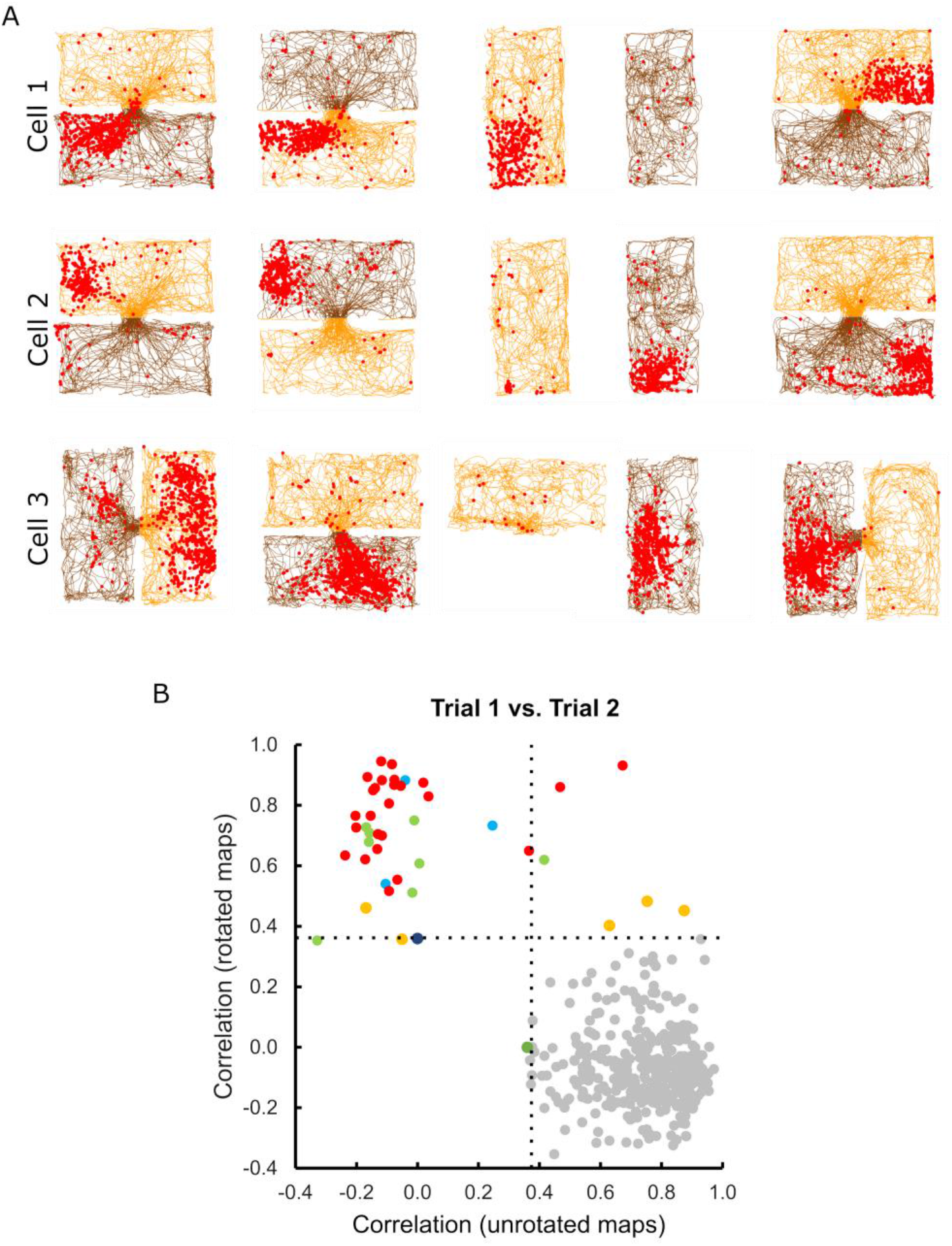
Odor switching. (A) Firing pattern of three example cells in which the firing location within a given box reversed its relationship to the box odors following rotations of 180ndegrees (top two cells) or 90 degrees (bottom cell). (B) Correlations between rate maps in the first trial vs. the second trial, made either with the maps in their canonical orientations (unrotated maps) or with one of the maps reversed 180 degrees (rotated maps). For the majority of fields (grey circles) the correlations were high in the unrotated condition and low in the rotated condition, meaning that the fields maintained the original relationship to the box configuration. For some fields (colored circles) the relationship reversed 180 degrees in the second trial such that the correlation was low in the unrotated condition and high in the rotated condition. These fields are color-coded by session; note that the rotated fields generally belonged to the same session, meaning that the rotation was an ensemble behavior.

To detect odor-switching systematically, we correlated every pair of rate maps for the baseline 1 with the rotation trial, in either the same orientation or with a 180° rotation: we then compared the result to a 95^th^ percentile threshold determined using a null distribution. When these two sets of correlations were plotted against each other (Figure 6B) then most data points, as expected, fell within the lower right quadrant, meaning there was a high correlation for the unrotated maps and low correlation when one of the maps was rotated 180 degrees. However a smaller but distinct cluster showed the reverse, occupying the upper left quadrant. Cells that showed this effect co-occurred, and never co-occurred with those that did not, indicating rotation of the whole ensemble. This effect was more evident at the beginning of the session and had disappeared by the last trial (data not shown), suggesting that the system had consolidated the relationship between odor and direction (see Discussion).

## Discussion

In order to investigate how place cells resolve environmental ambiguities, we recorded the cells in a two-compartment environment, the ‘2-box’, which has several symmetries that are broken with experimenter-controlled cues. Global rotational symmetry is broken with odor (assisted by using a constant start-box, during initial learning), while local rotational symmetry is broken with the within-box layout (the cue card and the doorway). The experiment was motivated by the recent finding that head direction (HD) cells in dysgranular retrosplenial cortex (dRSC) show two types of responding in this environment, with some cells sensitive to the local and some to the global cues (Jacob et al., 2017). We wanted to see if place cells, connected to RSC, would show this mixed pattern, expressing information about the ambiguous cues that could be useful for cognitive mapping.

We looked for three possible response patterns (Figure 1), but found only two of them. Specifically, we found that some place cells repeated their firing fields between the boxes, while some altered their firing altogether (remapped). However, we saw virtually no cells that rotated their firing to follow the rotated layouts. It thus seems that the directional ambiguity seen in RSC has been resolved by the time information reaches the place cells. Importantly, an additional observation was that although the orientation of the firing-fields ensemble generally matched that of the odor cues, sometimes the whole pattern dissociated from them and rotated *en bloc*. And finally, we observed that some types of symmetry-breaking were complete (affected all the cells together) and some partial.

Three things were notable about the above observations. First, the odor cues, which have no inherent directionality, were able to orient the place fields, even alone (in the single-box trials). Second, we had expected to see partial rotational remapping, whereby some cells might rotate their fields to follow the rotated environment cues. This is because the directionally ambiguous signal found in dRSC could potentially have found its way into the place cell system via the RSC-HPC interconnections; and also because a previous experiment dissociating directional and local cues did find that a subset of CA1 cells would follow the local cues (Knierim, 2002). Our observation is, by contrast, more similar to that of Fuhs et al. (2005) who found remapping but no rotation (albeit in a small sample from only two rats). It seems that no trace of the directional ambiguity remains in the place cell map.

The final notable observation from the present study was that the place field ensemble could dissociate from the odor cues without remapping (Figure 6). The reason this is surprising is that the cells definitely could respond to the odor cues, because the odors could orient the ensemble firing pattern, including (as mentioned) in the single-box trials. We had thus assumed that the remapping cells were also using the odors to distinguish the boxes and position their fields appropriately, since odors have been previously shown to influence such positioning (Anderson & Jeffery, 2003). However, that the ensemble would sometimes decouple from the odors and rotate *en bloc* suggests that odor-induced remapping was not always at play here.

How, then, did the remapping cells distinguish the compartments in order to remap, rather than rotate? The answer might lie with the only other symmetry-breaking information available, which is the relationship of the visual scene, arising from the layout, to global direction. If we suppose that in the lemon compartment the cue card is ‘North’ and the doorway ‘East’, then in the vanilla compartment the cue card is ‘South’ and the doorway ‘West’. Cells could thus, in principle, distinguish the compartments if they could associate these scenes with the global directional signal, conditional on the current odor. This notion is unpacked, below.

### Boundary vectors and context gating

An association between global direction and local environmental cues was earlier proposed by Burgess and colleagues, to explain why place fields ‘follow’ environment walls when these are moved relative to each other (Barry et al., 2006; Hartley, Burgess, Lever, Cacucci, & O’Keefe, 2000; John O’Keefe & Burgess, 1996). In their ‘boundary vector’ (BV) model (Figure 7A), each place cell is driven by inputs from some (not all) of the walls located at given allocentric directions. The BV model can explain the behavior of the cells that repeated their fields, because there are two ‘East’ and two ‘West’ walls in the 2-box.

**Figure 7.**
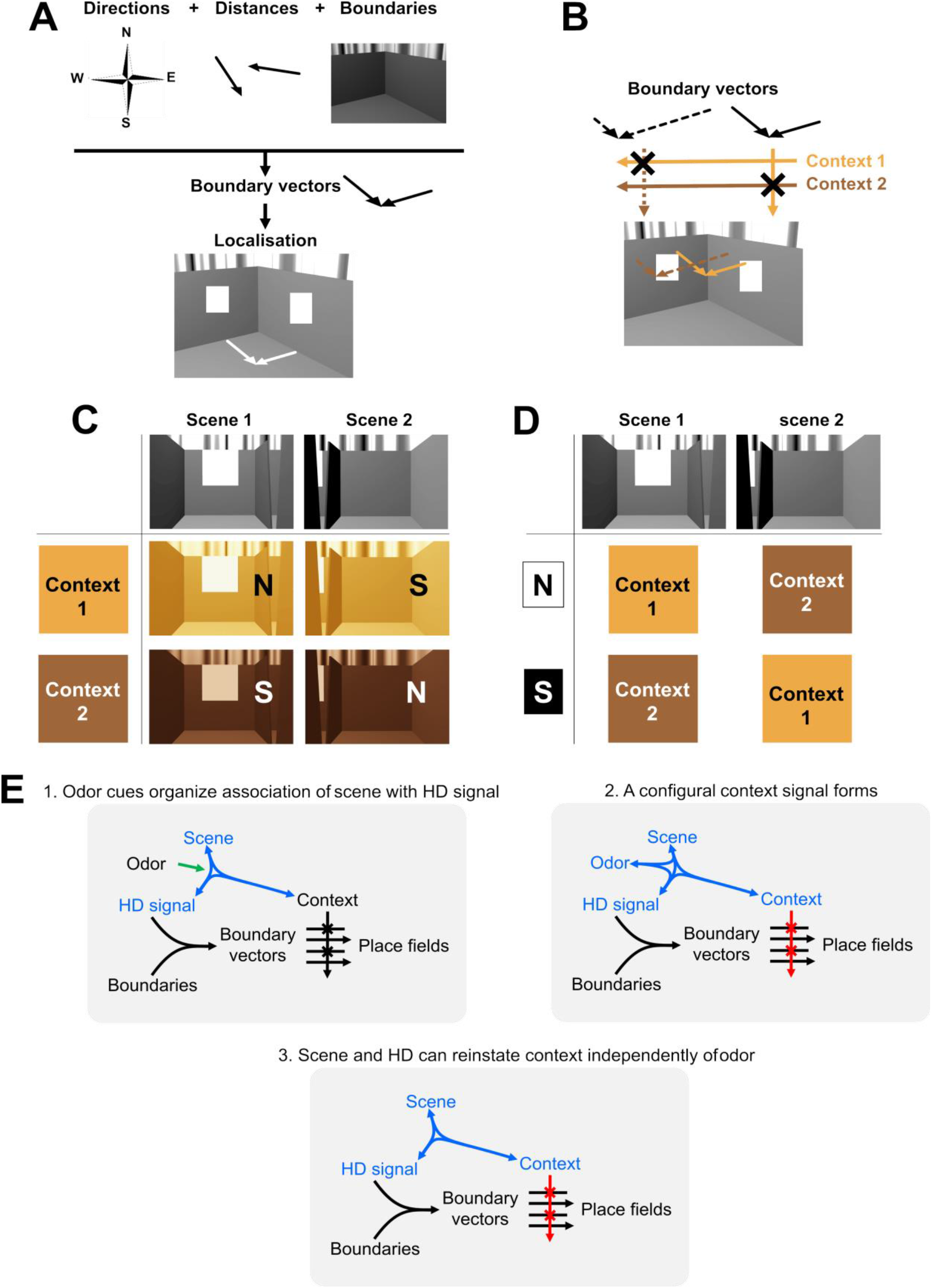
The logic of the configural encoding of sensory inputs driving place cells. (A) The boundary vector (BV) model of Burgess and colleagues (Barry et al., 2006). Separate signals about facing direction, boundaries and the distance from the boundaries combine to form boundary vectors (signals conveying distance to boundaries located at a given direction). When two or more orthogonal boundary vectors meet, they uniquely specify a location in the environment. (B) The context gating model of Jeffery et al. (Hayman & Jeffery, 2008; K J Jeffery et al., 2004). This extends the BV model by proposing that contextual cues control which boundary vectors can drive a place cell. (C) The configural combining of context and viewpoint to recover heading direction. In Context 1 (e.g., the lemon box in the 2-box) then the rat sees Viewpoint 1 when ‘North’ HD cells are active, and Viewpoint 2 when ‘South’ are active (S). In Context 2 the reverse holds. This configural signal (viewpoint + direction) can then input to boundary vectors. (D) The configural encoding of viewpoint and direction to recover context. A direction signal, which may have come from the process in (C) or may alternatively have come from spontaneous jumping of the HD network to the opposing orientation, combines with the viewpoint. If ‘North’ (N) HD cells are active when the rat sees Viewpoint 1 then it must be in Context 1, etc. This information can then be used to gate the boundary vectors driving the place cells, retrieving Context 1 or Context 2 pattern as appropriate. (E) Configural association of contextual cues with two organizational roles. 1. The odor cues, which are the only symmetry-breaking cues, organize the stable association of the visual scene with one set of HD cells. 2.The configural cue set (blue) generates a context signal that gates (red) the boundary inputs on to the place cells. 3. The context configuration can still work in the absence of some of its elements, explaining how the system can still use context to break the environment symmetry independently of the original symmetry-breaking cue.

The BV model does not explain all the results in the 2-box, however, because it predicts that every place cell should repeat its fields (except at the doorway) and thus does not account for the remapping cells. To explain remapping, such as when environment odor or color is changed, we have previously extended the BV model to include contextual cues (Jeffery et al., 2004; Hayman and Jeffery, 2008; Figure 7B). In our ‘contextual gating’ (CG) model, the subset of BVs that drives a place cell is selected (gated) by the extant context cues. The CG model can thus explain the remapping cells in the 2-box. The co-occurrence of remapping and repeating cells can further be explained by assuming a heterogeneity of inputs, in which repeating cells receive un-gated BV inputs while the remapping cells receive context-gated ones.

However, the CG model does not explain the results of the present experiment, in which we saw occasional decoupling of the place field ensemble from the odor context, because when this happens the BVs driving the remapping place cells have dissociated from their contexts, so how are these cells able to fire in their usual locations?

We come back to this issue in a moment. First, however, we consider the issue of how head direction is determined in the 2-box, given that this environment lacks a global polarizing visual cue.

### Environmental determination of the HD signal

The BV model and its CG variant assume the presence of a unitary global directional signal. Consistent with this, in our experiment we never saw cells rotate to follow the environment layout cues independently of the ensemble, so we assume that a unitary HD signal had been passed to all the place cells together. The question arises as to how this signal is aligned to the environment. In most experiments, directional information is provided to HD cells by a single polarizing cue such as a cue card on the wall of the recording chamber, but in our 2-box there are two such cards, in opposing directions. We broke this twofold structural symmetry by scenting the boxes with different odors: we know that HD cells are able to use this information, based on direct observation in RSC HD cells (Jacob et al., 2017) and also inferred here from the ensemble behavior of our place cells. But how does odor orient the HD signal, given odor’s inherent non-directionality? Also: given this role of odor in orienting the signal, how is it that the signal sometimes dissociates from the odors?

In fact, there *is* actually directional information provided by the odors, since the lemon and vanilla compartments are spatially separated and thus could in principle define an axis across the space. However the place cell patterns persisted even in the single compartments, so this axis computation seems unlikely. We instead propose an alternative explanation, which is that odor (or another equivalent non-spatial context signal) can drive the head direction signal by forming a configural association with the visual scene as perceived at a given facing direction (Figure 7C): that is, a three-way association between odor, head direction signal and layout (scene). Thus, if (for example) the ‘North’ HD cells were active when the rat was in the lemon box for the first time and facing the cue card, then the combination of lemon odor plus the cue card scene could associate with this subset of cells by Hebbian learning. Subsequently, the co-occurrence of the odor and that visual scene can re-activate the correct head direction cells. With the ‘South’ HD cells, although the visual scene is the same, the presence of the other (vanilla) odor forms a distinguishable configuration that prevents confusion.

Another way of thinking about this is to formulate it in the standard ring attractor framework within which theoretical discussions of HD cells are usually situated. By this view, the two identical cue card scenes drive the HD ring attractor at two different places 180 degrees apart, North and South; the odors break this ambiguity by selectively associating the cue card scene inputs to the ‘North’ or ‘South’ cells, as appropriate for that box, adding additional drive to one or other of the ring attractor locations and thus breaking the activation symmetry. This function for the odor cues is analogous to the gating role proposed for context in the CG model of place cell positioning.

### Two roles for context

The formation of a configural representation comprising odor, scene and HD signal, bound together, allows for ‘context’ to become a richer notion than simply the odor of a box. Because of the associations formed, the signal can persist even if one component transiently drops out. For example, if the rat closes its eyes (losing the scene inputs) it still knows which box it is in based on the directional and odor cues. If it becomes directionally disoriented (e.g. by an experimenter) it can retrieve heading based on the scene it can see plus the odor. If the odor signal is briefly lost, box identity can be retrieved using the combination of heading plus visual scene. This persistence of the context signal in the absence of one of its elements provides an explanation for why the whole ensemble sometimes rotates independently of the odor cues and yet the place cells retain their usual relative firing locations. It may also explain why place cells can sometimes generate independent representations for the same environment (Barnes, Suster, Shen, & McNaughton, 1997; Sheintuch et al., 2020).

This configural context signal has two separable roles. One is to define the orientation of the environment, as described above: the other is to interact with the BV inputs and control which ones drive the place cells (gating the BVs, as described earlier). This can explain how it is that the place field ensemble can act coherently in the directional domain (all cells orient together) but incoherently in the spatial domain (yielding both repeating and remapping cells).

### Anatomical locations of the interactions

We can speculate about the possible location of these processes based on the known anatomy and physiology of the region. Visual scene information from visual cortex may possibly pass through a scene-selective region such as parahippocampal cortex. Olfactory or other context information comes via the peripheral senses to a multimodal sensory area, which could be retrosplenial cortex or even hippocampus. Head direction information comes from an interaction of cortical and subcortical brain regions that combine self-motion information with static environmental information.

Where do these signals converge and interact? We propose that retrosplenial cortex is the most likely site. The Jacob et al (2017) experiment in the 2-box found cells that respond to visual layout independently of context (the so-called ‘bidirectional cells’), as well as directional cells that are context-sensitive (the classic HD cells). RSC receives context either directly from hippocampus or indirectly via the subiculum (see Vann et al., 2009; Mitchell et al., 2018 for review) and receive scene information from the visual cortex directly (Thomas Van Groen & Wyss, 2003; T van Groen & Wyss, 1992) or from as-yet-unidentified scene-selective regions of cortex. We propose here that this information can set the HD signal. This differs from previous formulation of HD cells in which that signal produces a unidirectional influence on place cells, and suggests that the construction of HD signals is, like many other signals in the brain, the product of feed-back as well as feed-forward inputs. Recordings from these areas as the cues are manipulated will be required to confirm the existence and determine the formation mechanism of these configural signals.

## Acknowledgements

This research was funded in part by the Wellcome Trust [Grant number WT103896AIA]. For the purpose of open access, the author has applied a CC BY public copyright licenses to any Author Accepted Manuscript version arising from this submission. The work was also supported by a grant from the Biotechnology and Biological Sciences Research Council (BB/Joo9792/1) to K.J.J, and an A*STAR (Singapore) National Science Scholarship to HYC.

## Author contributions

Han Cheng: Conceptualization, Methodology, Software, Investigation, Formal analysis, Data curation, Visualization, Writing – Original Draft, Review & Editing; Dorothy Overington: Conceptualization, Methodology, Software, Investigation, Formal analysis, Data curation; Kate Jeffery: Conceptualization, Methodology, Data curation, Supervision, Resources, Funding acquisition, Writing – Original Draft, Review & Editing.

The authors are grateful to Joanna Holeniewska for technical support during the project’s execution.

## Data and materials availability

Summarized datasets are documented in an online repository xxx, and the analysis code can be found on xxx (xxx).

## References

Anderson, M. I., & Jeffery, K. J. (2003). Heterogeneous modulation of place cell firing by changes in context. J.Neurosci, 23(26), 8827–8835.

Barnes, C. A., Suster, M. S., Shen, J., & McNaughton, B. L. (1997). Multistability of cognitive maps in the hippocampus of old rats. Nature, 388(6639), 272–275.

Barry, C., Lever, C., Hayman, R., Hartley, T., Burton, S., O’Keefe, J., … Burgess, N. (2006). The boundary vector cell model of place cell firing and spatial memory. Reviews in the Neurosciences, 17(1–2), 71–97.

Duvelle, É., Grieves, R. M., Hok, V., Poucet, B., Arleo, A., Jeffery, K. J., & Save, E. (2019). Insensitivity of place cells to the value of spatial goals in a two-choice flexible navigation task. Journal of Neuroscience, 39(13), 2522–2541. 10.1523/JNEUROSCI.1578-18.2018

Fuhs, M. C., VanRhoads, S. R., Casale, A. E., McNaughton, B., & Touretzky, D. S. (2005). Influence of path integration versus environmental orientation on place cell remapping between visually identical environments. Journal of Neurophysiology, 94(4), 2603–2616. 10.1152/jn.00132.2005

Grieves, R. M., Jenkins, B. W., Harland, B. C., Wood, E. R., & Dudchenko, P. A. (2016). Place field repetition and spatial learning in a multicompartment environment. Hippocampus, 26(1), 118–134. 10.1002/hipo.22496

Groen, Thomas Van, & Wyss, J. M. (2003). Connections of the retrosplenial granular b cortex in the rat. Journal of Comparative Neurology, 463(3), 249–263. 10.1002/cne.10757

Harland, B., Grieves, R. M., Bett, D., Stentiford, R., Wood, E. R., & Dudchenko, P. A. (2017). Lesions of the head direction cell system increase hippocampal place field repetition. Current Biology, 27(17), 2706–2712.e2. 10.1016/j.cub.2017.07.071

Hartley, T., Burgess, N., Lever, C., Cacucci, F., & O’Keefe, J. (2000). Modeling place fields in terms of the cortical inputs to the hippocampus. Hippocampus, Vol. 10, pp. 369–379.

Hayman, R. M., & Jeffery, K. J. (2008). How heterogeneous place cell responding arises from homogeneous grids - A contextual gating hypothesis. Hippocampus, 18(12). 10.1002/hipo.20513

Jacob, P.-Y., Casali, G., Spieser, L., Page, H., Overington, D., & Jeffery, K. (2017). An independent, landmark-dominated head-direction signal in dysgranular retrosplenial cortex. Nature Neuroscience, 20(2), 173–175. 10.1038/nn.4465

Jeffery, K.J. (2007). Integration of the sensory inputs to place cells: What, where, why, and how? Hippocampus, 17(9). 10.1002/hipo.20322

Jeffery, K J, Anderson, M. I., Hayman, R., & Chakraborty, S. (2004). A proposed architecture for the neural representation of spatial context. Neuroscience and Biobehavioral Reviews, 28(2). 10.1016/j.neubiorev.2003.12.002

Jeffery, K J, Donnett, J. G., Burgess, N., & O’Keefe, J. M. (1997). Directional control of hippocampal place fields. Experimental Brain Research, 117(1). 10.1007/s002210050206

Jeffery, Kate J. (2023). Symmetries and asymmetries in the neural encoding of 3D space. Philos. Trans. R. Soc. Lond. B Biol. Sci., 378(1869), 20210452.

Kadir, S. N., Goodman, D. F. M., & Harris, K. D. (2014). High-dimensional cluster analysis with the masked EM algorithm. Neural Computation, 26(11), 2379–2394. 10.1162/NECO_a_00661

Knierim, J. J. (2002). Dynamic interactions between local surface cues, distal landmarks, and intrinsic circuitry in hippocampal place cells. Journal of Neuroscience, 22(14), 6254–6264. 10.1523/jneurosci.22-14-06254.2002

Leutgeb, S., Leutgeb, J. K., Barnes, C. A., Moser, E. I., McNaughton, B. L., & Moser, M. B. (2005). Independent codes for spatial and episodic memory in hippocampal neuronal ensembles. Science, 309(5734), 619–623. 10.1126/science.1114037

Mamad, O., Stumpp, L., McNamara, H. M., Ramakrishnan, C., Deisseroth, K., Reilly, R. B., & Tsanov, M. (2017). Place field assembly distribution encodes preferred locations. In PLoS Biology (Vol. 15). 10.1371/journal.pbio.2002365

Mitchell, A. S., Czajkowski, R., Zhang, N., Jeffery, K. J., & Nelson, A. J. D. (2018). Retrosplenial cortex and its role in spatial cognition. Brain and Neuroscience Advances, 1, doi: 10.1177/2398212818757098.

Muller, R. U., & Kubie, J. L. (1987). The effects of changes in the environment on the spatial firing of hippocampal complex-spike cells. Journal of Neuroscience, 7(7), 1951–1968. 10.1523/jneurosci.07-07-01951.1987

O’Keefe, J, & Nadel, L. (1978). The Hippocampus as a Cognitive Map. Clarendon Press.

O’Keefe, John, & Burgess, N. (1996). Geometric determinants of the place fields of hippocampal neurons. Nature, 381(6581), 425–428. 10.1038/381425a0

Page, H. J. I., & Jeffery, K. J. (2018). Landmark-based updating of the head direction system by retrosplenial cortex: A computational model. Frontiers in Cellular Neuroscience, 12(July), 1–17. 10.3389/fncel.2018.00191

Sharp, P. E., Kubie, J. L., & Muller, R. U. (1990). Firing properties of hippocampal neurons in a visually symmetrical environment: contributions of multiple sensory cues and mnemonic processes. J.Neurosci., Vol. 10, pp. 3093–3105.

Sheintuch, L., Geva, N., Baumer, H., Rechavi, Y., Rubin, A., & Ziv, Y. (2020). Multiple maps of the same spatial context can stably coexist in the mouse hippocampus. Current Biology, 30(8), 1467–1476.

Skaggs, W. E., McNaughton, B., Gothard, K. M., & Markus, E. J. (1992). information theoretic approach to deciphering the Hippcampal code. Advances in Neural Information Processing Systems, 5, 1030–1038.

Spiers, H. J., Hayman, R. M. A., Jovalekic, A., Marozzi, E., & Jeffery, K. J. (2015). Place field repetition and purely local remapping in a multicompartment environment. Cerebral Cortex, 25(1), 10–25. 10.1093/cercor/bht198

Taube, J. S., & Burton, H. (1995). Head direction cell activity monitored in a novel environment and during a cue conflict situation. Journal of Neurophysiology, 74(5), 1953–1971.

Taube, J. S., Muller, R. U., & JB, J. R. (1990). Head-direction cells recorded from the postsubiculum in freely moving rats. II. Effects of environmental manipulations. Journal of Neuroscience, 10(2), 436–447.

Tolias, A. S., Ecker, A. S., Siapas, A. G., Hoenselaar, A., Keliris, G. A., & Logothetis, N. K. (2007). Recording chronically from the same neurons in awake, behaving primates. Journal of Neurophysiology, 98(6), 3780–3790. 10.1152/jn.00260.2007

van Groen, T, & Wyss, J. M. (1992). Connections of the retrosplenial dysgranular cortex in the rat. The Journal of Comparative Neurology, 315(2), 200–216. 10.1002/cne.903150207

Vann, S. D., Aggleton, J. P., & Maguire, E. A. (2009). What does the retrosplenial cortex do? Nature Reviews. Neuroscience, 10(11), 792–802. 10.1038/nrn2733

Zhang, N., Grieves, R. M., & Jeffery, K. J. (2022). Environment symmetry drives a multidirectional code in rat retrosplenial cortex. Journal of Neuroscience, 42(49), 9227–9241.

